# Sequence-based model of gap gene regulatory network

**DOI:** 10.1101/015776

**Authors:** Konstantin Kozlov, Vitaly Gursky, Ivan Kulakovskiy, Maria Samsonova

## Abstract

**Background:** The detailed analysis of transcriptional regulation is crucially important for understanding biological processes. The gap gene network in *Drosophila* attracts large interest among researches studying mechanisms of transcriptional regulation. It implements the most upstream regulatory layer of the segmentation gene network. The knowledge of molecular mechanisms involved in gap gene regulation is far less complete than that of genetics of the system. Mathematical modeling goes beyond insights gained by genetics and molecular approaches. It allows us to reconstruct wild-type gene expression patterns *in silico*, infer underlying regulatory mechanism and prove its sufficiency.

**Results:** We developed a new model that provides a dynamical description of gap gene regulatory systems, using detailed DNA-based information, as well as spatial transcription factor concentration data at varying time points. We showed that this model correctly reproduces gap gene expression patterns in wild type embryos and is able to predict gap expression patterns in *Kr* mutants and four reporter constructs. We used four-fold cross validation test and fitting to random dataset to validate the model and proof its sufficiency in data description. The identifiability analysis showed that most model parameters are well identifiable. We reconstructed the gap gene network topology and studied the impact of individual transcription factor binding sites on the model output. We measured this impact by calculating the site regulatory weight as a normalized difference between the residual sum of squares error for the set of all annotated sites and the set, from which the site of interest was left out.

**Conclusions:** The reconstructed topology of the gap gene network is in agreement with previous modeling results and data from literature. We showed that 1) the regulatory weights of transcription factor binding sites show very weak correlation with their PWM score; 2) sites with low regulatory weight are important for the model output; 3) functional important sites are not exclusively located in cis-regulatory elements, but are rather dispersed through regulatory region. It is of importance that some of the sites with high functional impact in *hb*, *Kr* and *kni* regulatory regions coincide with strong sites annotated and verified in Dnase I footprint assays.

## Background

The detailed analysis of transcriptional regulation is crucial for understanding biological processes, and interest in this problem grows due to new large-scale data acquisition techniques. However despite our expanding knowledge of the biochemistry of gene regulation, we lack a quantitative understanding of this process at a molecular level. We do not understand the mechanism of transcription factor (TF) interactions with adaptor proteins, basal transcriptional machinery and chromatin. We do not know why some cis-regulatory elements (CREs) are modular, while other are scattered over many kilobases of DNA. We cannot effectively predict the aspects of spatiotemporal expression mediated by a particular DNA region and which set of transcription factor binding sites (TFBS) forms a functional CRE.

The segment determination network in *Drosophila* attracts large interest among researches studying mechanisms of transcriptional regulation. The body of fruit fly consists of repeated morphological units called segments. The borders of segments are demarcated (determined) simultaneously during the blastoderm stage, just before the onset of gastrulation. The segment determination is under control of hierarchical cascade of segmentation genes, most of which are transcriptional regulators. These genes fall into 4 classes. At the bottom of the cascade are the maternal co-ordinate genes *bicoid* (*bcd*, one letter code – B) and *caudal* (*cad*, one letter code – C). The other groups of genes are gap genes (*Kruppel* (*Kr*, one letter code – K), *giant* (*gt*, one letter code – G), *hunchback* (*hb*, one letter code – H), *knirps* (*kni*, one letter code – N), *tailess* (*tll*, one letter code – T) and *huckebain* (*hkb*, one letter code – J), pair-rule and segment-polarity genes.

There is a large amount of experimental data available about the segment determination system. The gap gene system implements the most upstream regulatory layer of the segmentation gene network. It receives inputs from long-range protein gradients encoded by maternal coordinate genes and establishes discrete territories of gene expression. In this process the gap gene cross-regulation plays important role. The formation of gap gene expression domains is a dynamic process: the domains do not form in one place, but instead in the posterior half of the embryo they shift anteriorly during cleavage cycle 14.

At the molecular level we know the genomic location of many functionally verified CREs, as well as identity and binding affinity of sites for relevant regulating TFs. A wealth of genome scale functional studies provide data on Chip-Seq, RNA-Seq and DNasel accessibility measurements. The analysis of these datasets demonstrated that maternal co-ordinate and gap TFs bind to thousands of sites across the *Drosophila* genome and that the dominant force in cells that modifies the intrinsic DNA specificity of TFs is the inhibition of DNA binding by chromatin [1]. High resolution imaging and image processing techniques provide spatiotemporal data on segmentation gene expression at cellular resolution [2].

In spite of these efforts we still do not understand the molecular mechanisms involved in gap gene regulation. It is known that the the gap regulatory regions usually contain several CREs driving expression in a precise spatiotemporal pattern and often containing large number of apparent redundant sites for the same TF. Certainly this molecular complexity is important, however the mechanisms underlying it remain elusive.

Mathematical modeling extends the boundaries of genetics and molecular approaches. In general the sufficiency of inferred regulatory mechanism cannot be proven without reconstructing the system *ab initio*. Currently there is no assay, which accurately reproduces eukaryotic transcription *in vitro* from well-defined reagents. Mathematical modeling allows us to reconstruct wild type gene expression patterns *in silico*, to infer underlying regulatory mechanism and prove its sufficiency. Three major classes of mathematical models have been applied to model regulation in gap gene network: Boolean, differential equation-based and thermodynamic (also termed fractional occupancy) models [3].

Boolean models represent regulatory relations as logic gates and in the gap gene system they were applied to identify feedback loops which account for topology of gene network at steady-state.

The differential equation based models represent a regulatory network by differential equations, in which a set of molecules such as mRNAs and proteins interact by explicit rules defined in terms of kinetic equations. When applied to the gap gene system these models were able to infer regulatory interactions responsible for formation of the expression domain boundaries, as well as to explain mechanisms for the dynamic anterior shifts of gap domains. It should be noted that the deciphering of the mechanisms of domain motion would be impossible with classic genetic approaches in default of mutants defective for any specific domain shift.

Thermodynamic models rely on simple biophysical descriptions of DNA-protein interactions and statistical physics. They attempt to infer information about gene regulation from the sequences of CREs and the binding affinities of TFs to these elements. This formalism was used to model expression levels in constructs driving reporter gene expression from different gap gene regulatory elements.

It should be noted that all these models have advantages and limitations from the perspective of input data quantity, degree of complexity, and the time interval in which they can model gene expression. Boolean models are suitable to work with large amounts of data produced by genome-wide experiments, but they do not in general consider DNA sequence information. Thermodynamic-based models specifically take into account the features of CREs. However these models provide output for a particular time moment and do not capture the system dynamics. On the contrary differential equation models allow scientists to consider transcriptional regulation over continuous time intervals. The primary limitation of these models is the size of gene network, as the number of parameters rapidly grows with increase of gene number and the problem becomes computationally infeasible. Besides, the differential equation based models usually describe gene interactions in terms of activation/repression and the fine details of transcriptional regulation that thermodynamic-based models offer, are not included.

Evidently, to decipher the molecular mechanisms involved in gap gene regulation we need to understand how genetic information encoded in regulatory elements of these genes is translated into dynamical aspects of gap gene expression. This can be achieved by combining strength of both thermodynamic and differential equation based formalisms. Here we present a new model that provides a dynamical description of gap gene regulatory systems, using detailed DNA-based information, as well as spatial TF concentration data at varying time points. We showed that this model correctly reproduced gap gene expression patterns in wild type embryos and is able to predict gap expression patterns in mutants and reporter constructs.

## Results and discussion

### Sequence based model of gap gene network

We developed a new model of the gap gene regulatory network which takes as input the affinities of predicted TFBS together with spatial TF concentration data. The output of the model are spatial and temporal patterns of four gap genes *hb*, *Kr*, *gt*, and *kni* in the form of protein concentration profiles over about one hour of development.

The binding sites for TFs Bcd, Cad, Hb, Gt, Kr, Kni, Tll and Hkb were predicted using positional weight matrices (PWMs, see Additional file 1 and Methods). The predicted TFBS affinities were calculated based on the PWM score of the corresponding strongest site as in [4]. The spatial TF concentration data were taken from FlyEx database, which contains data on segmentation gene expression at cycles 13 and 14A of the early embryo development [5].

Our model consists of two layers. The first layer is a thermodynamic based calculation of the gene activation level. We adopt and modify a method of this calculation presented in [4]. The probability of transcriptional gene activation is assumed to be dependent on the rate of basal transcriptional machinery (BTM) recruitment, which is determined by different probabilities of all possible occupancy states of the regulatory region. Each occupancy state represents a different TF binding configuration on the DNA sequence. As many CREs require mechanisms such as synergy, cooperativity, quenching, and direct repression for proper function [6, 7, 8, 9, 10] the model incorporates additional mechanistic features such as short range repression and homotypic cooperativity in transcription factor-DNA binding [11].

The short range repression, also known as quenching, is a mechanism by which repressors influence activators only if they are bound within a “short range” of the activator binding site [12, 13]. According to this mechanism, a bound repressor cannot interact with the basal complex, but instead leads to a new configuration of the enhancer where its neighborhood in the DNA sequence becomes forbidden to binding by any other TF [4].

One feature of the model which can be incompatible with the gap gene network is the fact that the type of regulatory action (activation or repression) and its strength for a given TF is the same for all target genes. Previous modeling and experimental results showed that this is not true for gap genes, which may simultaneously exhibit self-activation and repression for other gap genes [14]. Taking this into account, we modified the model to allow different regulatory actions for TFs depending on a target gene, as described in more details in the next section.

Following [4] we consider that transcriptional output is proportional to the probability of the BTM binding. To model the spatio-temporal dynamics of mRNA and protein synthesis in the early embryo we write two sets of the reaction-diffusion differential equations [15, 16, 17]. We added the delay parameter to account for the average time between events of transcription initiation and corresponding protein synthesis.

We modeled, in one dimension, a region of the blastoderm corresponding to the central midline of the embryo. We consider a time period of cleavage cycles 13 and 14A. Cleavage cycle 14A is about one hour long and is divided into 8 temporal classes (T1-T8) of 6.5 minutes each. The number of nuclei along the A-P axis is doubled when going from cl3 to cl4. The model was fitted to data on gap protein concentrations from the FlyEx database [5]. Optimization was carried out by differential evolution (DEEP) method [18, 19] to minimize the combined objective function. This function is a sum of the residual sum of squared differences between the model output and data, weighted pattern generating potential and a penalty term, which limits a growth of regulatory weights. The weighted pattern generating potential was proposed in [20] to account not only for the magnitude of difference between model and data, but also for the direction of change.

The model outputs with the score of combined optimization function below 350000 were inspected visually and the solutions, which fit the data without visual defects were selected. We obtained eleven similar solutions which produced calculated expression patterns that closely match the gap gene expression profiles in the wild type embryo (Figure1).

**Figure 1.**
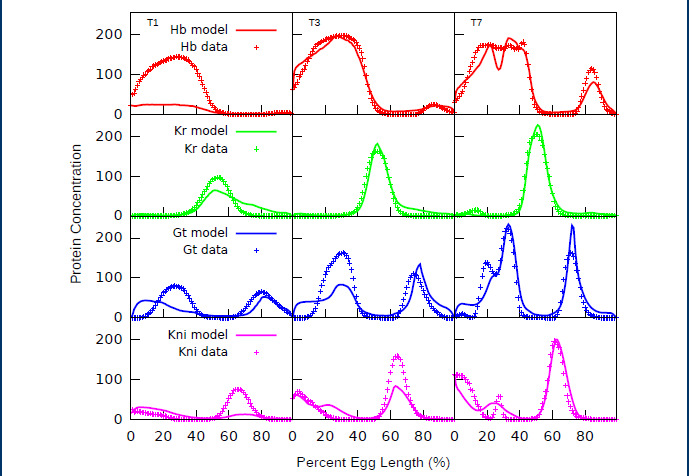
Model output for a representative network as compared to protein concentration profiles from the FlyEx database. Results are shown for 3 time moments - early (T1), middle (T3) and late (T7) cleavage cycle 14A. Though there are some defects in predicted patterns at T1, the model correctly reproduces the dynamic of the system.

To validate our fitting procedure we performed a four-fold cross-validation test. The entire dataset was partitioned randomly into four subsets. Then, the model was fitted using the data contained in three subsets (a training set). The obtained parameter values were used to make predictions for the subset left out (a test set) and the quality of prediction was estimated by calculation of the root mean square (*rms*) (see Methods section). This is repeated four times so that each subset is left out exactly once. This procedure resulted in the mean *rms* score 28.42 and standard deviation 1.29 that is comparable with the scores for original parameter sets *rms_mean_* = 27.15 and *rms_sd_* = 2.14. We applied Student’s t-test with Welch modification [21] to confirm that the difference between these *rms* scores is statistically insignificant, *P* > 0.10. Figure 2 shows the boxplot of the *rms* values for original and “cross-validation” networks.

**Figure 2.**
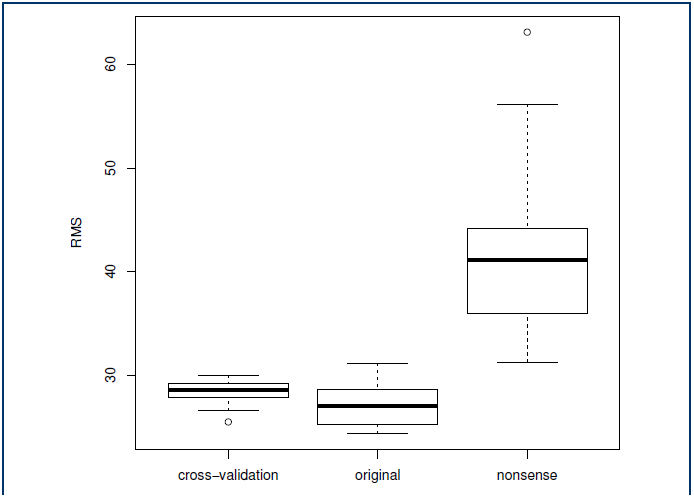
Box-and whisker plot of *rms* values obtained by fitting to biological data, in cross-validation test and by fitting to random dataset.

In order to further validate that the model is sufficient in data description we constructed a random dataset (“negative control”) in which the expression patterns of *kni* and *hb*, as well as expression patterns of *Kr* and gt were shuffled with respect to gene regulatory regions. Consequently, the data the model is fitted to may be considered “nonsense”. In this test we hoped that no parameter set could be found making the model output to coincide with “nonsense” data. We noted that a portion of resulting parameter sets has very small affinity constants (*K* < 10^−4^) for all TFBS of several TFs, and, hence, these TFs are almost switched off. Evidently, such a situation is not feasible and therefore we removed these parameter sets from further analysis. The mean *rms* score for the obtained set of parameter vectors was 41.07. The boxplot of the *rms* scores for biological and negative control data is presented in Figure 2. According to Student’s t-test with Welch modification *t* = 11.26, *P* — *value* = 5.101 × 10^−15^, consequently, the difference in *rms* mean values is statistically significant.

### Gene network topology

In segmentation network a TF can function as both activator and repressor. To account for the possibility of dual regulation we introduced the genetic inter-connectivity matrix *T^ab^*, which characterizes the action of TF *b* on gene *a*. The positive elements of the matrix are statistical weights *α_a_* of interaction between bound TF and the BTM, while negative elements correspond to the repressor strength *β_r_.* We assume that a bound repressor *R* acts via the short-range repression mechanism. We describe the topology of regulatory network by assigning the elements of *T* matrix into two categories: ‘activation’ (positive parameter values) and ‘repression’ (negative parameter values). The predicted topology corresponds to categories containing most of the parameter values (Table 1). The main features of the gap gene network topology are in agreement with previous modeling results and data from literature [14]. Bcd and Cad activate zygotic gap gene expression in a majority of circuits. Genes *hb*, *Kr*, *kni*, and *gt* exhibit autoactivation. Third, the reciprocal interactions between the trunk gap genes *Kr*, *hb*, *kni* and *gt* are repressive. An exception is activation of *hb* by Gt and Kr. Tll represses *Kr* and *gt*, and acts as activator of *hb* and repressor of *gt* in a majority of networks. For a majority of parameter sets Hkb represses *hb*, *Kr* and *gt*, but acts as *kni* activator in a half of networks.

**Table 1.** Prediction of network topology based on classification of T matrix elements. Numbers in cell define in how many networks a given interaction was classified as activation or repression. Columns correspond to TFs, rows to target genes.

| Name | *hb* | *Kr* | *gt* | *kni* | *bcd* | *cad* | *tll* | *hkb* |
| --- | --- | --- | --- | --- | --- | --- | --- | --- |
| *hb* | (11,0) | (1,10) | (3,8) | (0,11) | (11,0) | (8,3) | (7,4) | (3,8) |
| *Kr* | (9,2) | (10,1) | (0,11) | (1,10) | (4,7) | (8,3) | (0,11) | (1,10) |
| *gt* | (1,10) | (0,11) | (11,0) | (1,10) | (9,2) | (6,5) | (1,10) | (4,7) |
| *kni* | (0,11) | (0,11) | (0,11) | (11,0) | (11,0) | (8,3) | (4,7) | (5,6) |

### Parameter identifiability

For further studies we selected one of the parameter sets based on its best visual coincidence with experimental data and low *rms* value equal to 25.18. In this network (Table 2) *Kr* is activated by Bcd and slightly repressed by Hb. Cad activates *hb*, *Kr*, *gt*, but slightly represses *kni*. Tll activates *hb* and represses all other trunk gap genes. Hkb acts as a repressor. TFs Hb, Kr, Gt and Tll have high cooperativity constants *ω* close or equal to 5. On the other hand, Bcd and Cad received low cooperativity values close to 1 together with Kni and Hkb. Affinity binding constants *K* for a TF strongest site vary by three orders of magnitude between 0.0001347 for Hkb and 0.049862 for Kni.

**Table 2.** The parameter estimates for a representative network. Columns correspond to TFs, rows to target genes. K and ω are affinity and cooperativity constants respectively. Poorly identifiable interactions are marked with *

| Name | *hb* | *Kr* | *gt* | *kni* | *bcd* | *cad* | *tll* | *hkb* |
| --- | --- | --- | --- | --- | --- | --- | --- | --- |
| *hb* | 2571.86 | -10.96 | 472.97 | -703.89 | 3821.42 | 1232.89 | 824.14* | -5487.58 |
| *Kr* | 136.22 | 222.92* | -311.29 | -60.24 | 33.35 | 9084.28* | -3388.59 | -993.95* |
| *gt* | -1528.17 | -4250.83 | 8310.29 | -1513.01 | 918.23 | 2272.27 | -3612.94 | -1208.58 |
| *kni* | -3946.83 | -2424.43 | -3399.80 | 8973.61 | 5232.66 | -11.30 | -5126.47 | -9175.97 |
| $K$ | 0.005731 | 0.004891 | 0.036382 | 0.049862 | 0.008036 | 0.005595* | 0.000223 | 0.001347 |
| $\omega$ | 5.000000 | 4.958060 | 5.000000 | 1.000053 | 1.000012 | 1.000001* | 4.565639 | 1.132316* |

To understand how reliable our model is we performed the identifiability analysis of the model parameters estimated by fitting to experimental data.

We decide about the sensitivity of the model solution to parameter changes by calculating the confidence intervals for the estimated parameter values (see Methods). This calculation is performed under the assumption that error in data is normally distributed. The error in the gene expression data almost linearly increases with the mean concentration, as it happens for the Poisson distribution. We apply the variance-stabilizing transform 
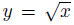
 to both data and model solution in order to make the error independent of the mean. The parameter estimates found for original objective turned out to be also the minimizers for the transformed one.

The predicted topology of regulatory network is based on the sign of the *T* matrix elements. We constructed confidence intervals for the parameter set from Table 2 in the vicinity of the model solution. Some values of regulatory parameters are small, and it is necessary to inspect the significance of the values or their signs. We classify parameters as non-identifiable if their confidence interval includes both positive and negative values and hence contains zero. It can be seen in Figure 3 that the non-identifiable regulatory parameters are autoregulation of *Kr* and the regulation of *Hb* by Tll, which means that we cannot make significant conclusions about these interactions. The regulatory parameters which involve Hkb as a repressor have large confidence interval. The same is true for the regulatory parameter which characterizes the action of Gt on *tll*. The analysis shows good identifiability of all other regulatory parameters. Therefore, the identifiability analysis sustains the gene network topology deduced from classification of parameter values only.

**Figure 3.**
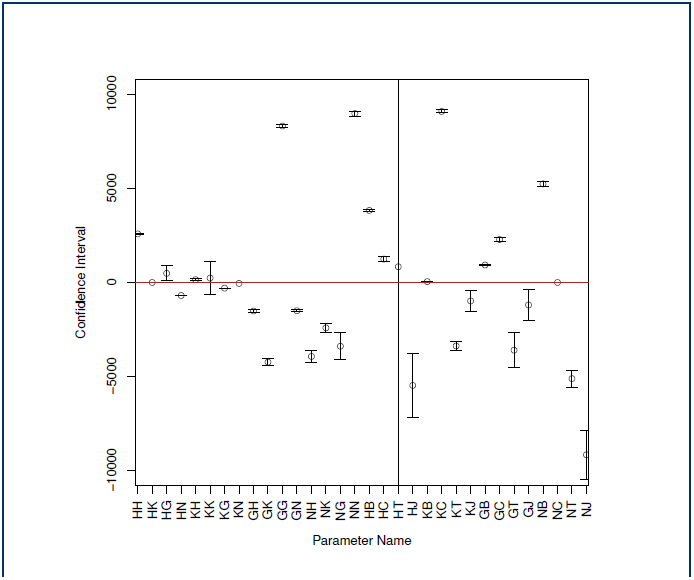
95% confidence intervals for estimates of the T matrix elements of a representative network. The parameter estimates are labeled by single-letter notations of genes: *hb*(H), *Kr*(K), *gt*(G), *kni*(N), *bcd*(B), *cad*(C), *tll*(T), *J*(HKb). The first letter corresponds to the target gene (e.g., HT stands for *T^HT^*). **HT** has the largest interval and the interval for **KK** crosses the zero axis.

The confidence intervals for thermodynamic parameters are given in Table 3. For most of these parameters the confidence intervals are small. The exceptions are cooperativity constants *ω* for Kr, Tll and Hkb, which have very large confidence intervals.

**Table 3.** Estimates and 95% confidence intervals for affinity and cooperativity constants *K* and ω in a representative network. Left and right interval borders are presented in columns marked “a” and “b” respectively.

| Parameter | Value | a | b |
| --- | --- | --- | --- |
| $K^H$ | 0.005731 | 5.663804e-03 | 5.798196e-03 |
| $K^K$ | 0.004891 | 4.705545e-03 | 5.076455e-03 |
| $K^G$ | 0.036382 | 3.604491e-02 | 3.671909e-02 |
| $K^N$ | 0.049862 | 4.933258e-02 | 5.039142e-02 |
| $K^B$ | 0.008036 | 7.944617e-03 | 8.127383e-03 |
| $K^C$ | 0.005595 | 5.547746e-03 | 5.642254e-03 |
| $K^T$ | 0.000223 | 2.106789e-04 | 2.353211e-04 |
| $K^J$ | 0.001347 | 1.180521e-03 | 1.513479e-03 |
| $\omega^H$ | 5.000000 | 4.829196e+00 | 5.170804e+00 |
| $\omega^K$ | 4.958060 | -4.610752e+00 | 1.452687e+01 |
| $\omega^G$ | 5.000000 | 4.499104e+00 | 5.500896e+00 |
| $\omega^N$ | 1.000053 | 7.967040e-01 | 1.203402e+00 |
| $\omega^B$ | 1.000012 | 9.202935e-01 | 1.079730e+00 |
| $\omega^C$ | 1.000001 | 9.090420e-01 | 1.090960e+00 |
| $\omega^T$ | 4.565639 | 7.928902e-02 | 9.051989e+00 |
| $\omega^J$ | 1.132316 | -2.311281e+01 | 2.537744e+01 |

The confidence intervals provide the full information about the parameter estimates only in case of parameter independency, otherwise the intervals are overestimated. Moreover, strong correlation between parameters may lead to their non-identifiability, because a change in one parameter value can be compensated by the appropriate changes of another parameters and, hence, does not significantly influence the solution. It was reported that parameters in the thermodynamic models, for example, affinity constants and cooperativity constants, may be correlated [22]. Because of that we investigated the dependencies between parameters using the collinearity analysis of the sensitivity matrix. This method allows to reveal correlated and hence non-identifiable subsets of parameters.

The sensitivity matrix was analyzed in the vicinity of the point in the parameter space that define the optimal model solution as described in the Methods section. The collinearity index *γ_κ_* (3) was computed for all the subsets of dimension *k* of the parameter set with the threshold value fixed at 4. For *k* = 3, this threshold value corresponded to approximately 90% pairwise Pearson correlation between columns of the sensitivity matrix. We identified poorly identifiable by finding 2-and 3-dimensional subsets with the collinearity index exceeding the threshold value (Table 4). It turned out that almost all parameter combinations in these subsets involve parameters defined as non-identifiable by exploration of the confidence intervals, namely regulatory parameter *T^KK^* for *Kr* autoactivation, regulatory parameter *T^KJ^*, which involves Hkb as a repressor and *Kr* as a traget gene, or cooperativity constant *ω_Hkb_* The correlation between parameters in this approach is related to large confidence intervals of parameter estimates. For example, very large confidence interval for both parameters *T^HT^* and *ω_Hkb_* can be explained by 52% correlation between these parameters. In the same way 93% correlation between *T^KK^* and *T^KJ^* explains large confidence intervals for these parameters.

**Table 4.**
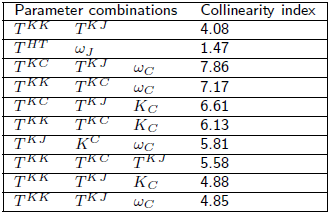
Two- and three-dimensional subsets of T-matrix elements with collinearity indices higher than 4. The collinearity index for the parameter combination *T^HT^* and *ω_Hkb_* does not exceed threshold, however these parameters show high correlation (Pearson correlation coefficient 52%) that explains their large 95% confidence intervals.

| Parameter combinations | Collinearity index |
| --- | --- |
| $T^{KK}$ $T^{KJ}$ | 4.08 |
| $T^{HT}$ $\omega_J$ | 1.47 |
| $T^{KC}$ $T^{KJ}$ $\omega_C$ | 7.86 |
| $T^{KK}$ $T^{KC}$ $\omega_C$ | 7.17 |
| $T^{KC}$ $T^{KJ}$ $K_C$ | 6.61 |
| $T^{KK}$ $T^{KC}$ $K_C$ | 6.13 |
| $T^{KJ}$ $K^C$ $\omega_C$ | 5.81 |
| $T^{KK}$ $T^{KC}$ $T^{KJ}$ | 5.58 |
| $T^{KK}$ $T^{KJ}$ $K_C$ | 4.88 |
| $T^{KK}$ $T^{KJ}$ $\omega_C$ | 4.85 |

It should be noted that the gene network topology revealed in this work is to a large extent in agreement with experimental evidences [14], however several disparities exist. In our model Bcd activates *Kr* in some networks and represses in the others. It was shown that in *bcd* mutant mothers *Kr* expression is not reduced but expands anteriorly [23]. This fact leads to proposal that high concentrations of Bcd repress *Kr* [23, 24], however this effect was later explained by the absence of the anterior *gt* and *hb* domains [25]. The activating effect of Bcd on *Kr* is supported by the fact that *Kr* expression in reporter constructs is activated by Bcd [26, 27]. The finding that *Kr* is still expressed in embryos from *bcd* mutant mothers has been explained by general transcription factor activation [28] or low levels of Hb [24, 29]. Our analysis does not allow us to make the unambiguous interpretation of the mechanisms of Hkb, Tll and Cad action as these TFs repress and activate target genes in much the same number of networks. It is believed that high concentrations of Cad at the posterior of the embryo activate gap genes. However at about 10 – 15 minutes before gastrulation Cad expression domain refines into a narrow posterior stripe [2]. The posterior *hb* domain is completely absent in *tll* mutants [30, 31], that suggests activation of posterior *hb* by Tll. Some data indicates that Hkb does repress *hb*, *Kr* and *gt*. For example, in *hkb* mutant embryos the posterior *hb* domain is unable to retract from the posterior pole [32]. Besides, in embryos mutant for the maternal gene *vasa* (*vas*), *tll* and *hkb* the *Kr* domain expands further posterior than in those mutant for *vas* and *tll* alone [33]. Finally, in embryos mutant for *tll* the posterior domain of *gt* expands less to the posterior pole that in *tll hkb* double mutants [34]. An explanation for the model failure to provide unambiguous prediction of the mechanism of Cad, Tll and Hkb action can be found in our analysis of parameter identifiability. This analysis showed that many parameters defining gap gene regulation by Hkb, Cad and Tll are non-identifiable (see Table 3, Table 4 and Figure 3) and therefore we cannot draw any conclusion about these interactions.

### Prediction of gap expression in Kr mutants and reporter constructs

We use parameters estimated on wild-type expression data to predict *in silico* gap gene expression patterns in *Kr* mutants and reporter constructs.

To simulate *Kr* null mutants we set the maximum synthesis rates 
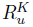
 and 
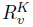
 for *Kr* to zero and fed the concentration profiles of TFs from mutant embryos to the model. Null mutation in *Kr* leads to significant decrease in gap gene expression levels in cycle 14A. Also, the posterior Gt domain exhibits a large shift, and positions of posterior Gt and Kni domains overlap [17]. Our model reproduces these features correctly: posterior Gt domain shifts anteriorly and coincides with abdominal Kni domain and the expression levels of gap genes *hb, gt*, and *kni* are reduced (Figure 4).

**Figure 4.**
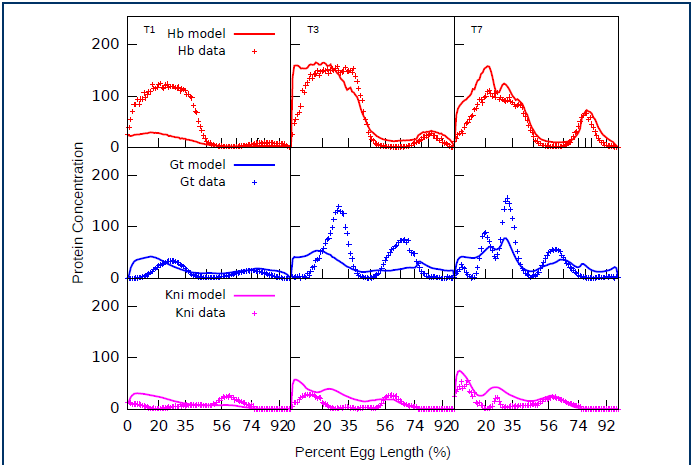
In silico predictions of gap gene expression patterns in *Kr*^−^ mutants. Parameters were fitted using wild type data only. The model correctly reproduces the characteristic features of gap gene expression in mutants, namely the decrease of gap gene expression levels and the anterior shift of *gt* domain.

To model gap gene expression driven by reporter constructs we take as input only those TFBS that overlap with CRE contained in a reporter. The CRE coordinates were taken from RedFly database [35]. The following reporter constructs were used: *gt*_*(-3)*, *Kr*_*CD1*,*Kr*_*730*, *kni*_*223+64* and *kni*_*kd* . The *gt*_*(-3)* construct contains CRE that drives the reporter gene expression in the *gt* posterior domain, *kni*_*kd* contains CRE that reproduces *kni* posterior expression and both *Kr*_*CD1* and *Kr*_*730* are expressed in the central *Kr* domain [26, 36, 35]. The *kni*_*223+64* construct contains CRE that conducts the posterior *kni* expression [37]. As is evident from Figure 5 the model is able to correctly predict the spatial features of expression in all constructs: the positions of predicted expression patterns coincide well with the positions of expression domains in constructs, as well as with the positions of corresponding gap gene endogenous domains. It should be also noted that enzymatic qualitative method used for staining precludes the comparison of expression levels predicted *in silico* and driven by constructs.

**Figure 5.**
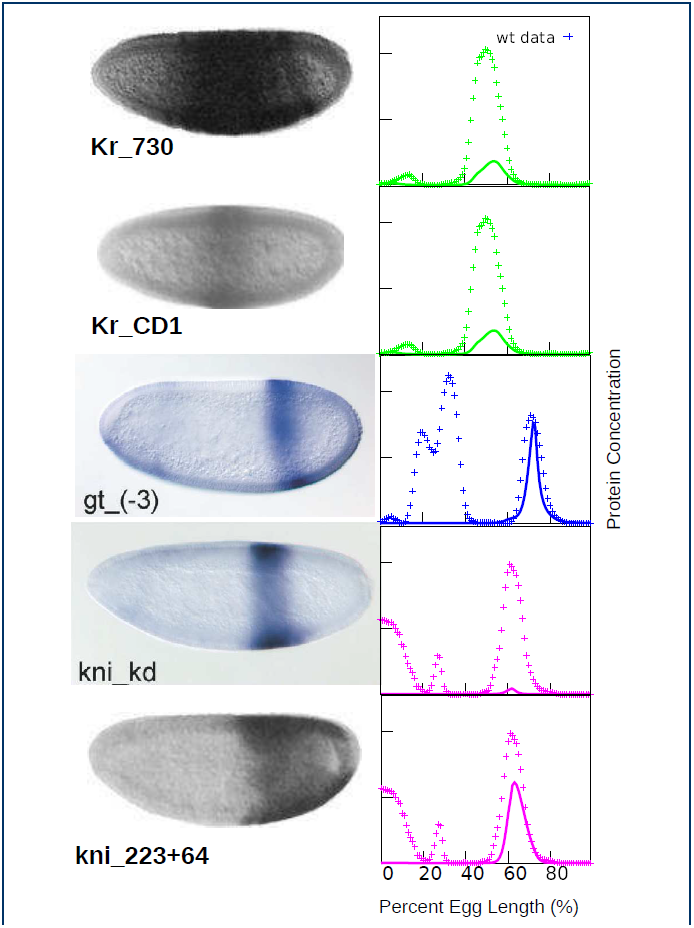
In silico prediction of gap gene expression patterns in reporter constructs. The construct *gt*_*(-3)* contains CRE that drives the reporter gene expression in the *gt* posterior domain, *kni*_*kd* contains CRE that reproduces *kni* posterior expression, both *Kr_730* and *Kr*_*CD1* are expressed in the *Kr* central domain. Both *kni*_*223*+*64* and *kni*_*kd* constructs contain CRE that conducts the posterior *kni* expression [37].

These results convincingly demonstrate that our model is able to correctly predict expression patterns in null mutants and reporter constructs from fits to wild-type data only. This provides an independent proof of model correctness and opens a way for its application for deciphering the mechanisms of transcriptional regulation and gene expression, as will be discussed below.

### Contribution of individual TFBS to gap gene expression

Functional genomics studies of animal regulatory regions lead to the hypothesis that transcription factors bind to a majority of genes over a quantitative series of DNA occupancy levels and that the weak regulatory interactions may be of biological significance [38]. Here we use our model to corroborate this idea. Specifically we tried to find the answer on three questions. Are TFBS of small functional impact still important for the model output? Does the correlation between the functional significance of TFBS and its binding affinity exist? Are functional important sites dispersed through regulatory region or predominately located within CREs?

To estimate the functional impact of an individual TFBS, i.e. its influence on the overall quality of model output, we define the regulatory weight (RW) of TFBS *w_r_* as

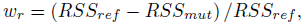

where *RSS_ref_* is the residual sum of squares error between the wild type expression data and the model solution for the full set of annotated sites, and *RSS_mut_* is the same quantity calculated with the site of interest excluded.

We have calculated the RW for each annotated site and for each gap gene regulatory region. In bioinformatics PWM models are generally used to calculate the BS affinity. We found that the RWs of TFBS show very weak correlation with their PWM score (Spearman’s rank correlation coefficient *ρ* = 0.15, *P* = 3.5 × 10^−6^; Pearson correlation coefficient *r* = 0.17, *P* = 2.7 × 10^−7^). This suggests that the influence of a TFBS on the phenotype is to a great extent explained not by the binding strength *per se* but by the way the binding sites are involved in the gene regulatory network.

In Figure 6 we plot the RW of TFBS relative to their position in a regulatory region. Some sites overlap with the reporter construct CREs, while the other do not. A number of sites from both these categories have high impact on the model solution, however the majority of sites have relatively low individual impact.

**Figure 6.**
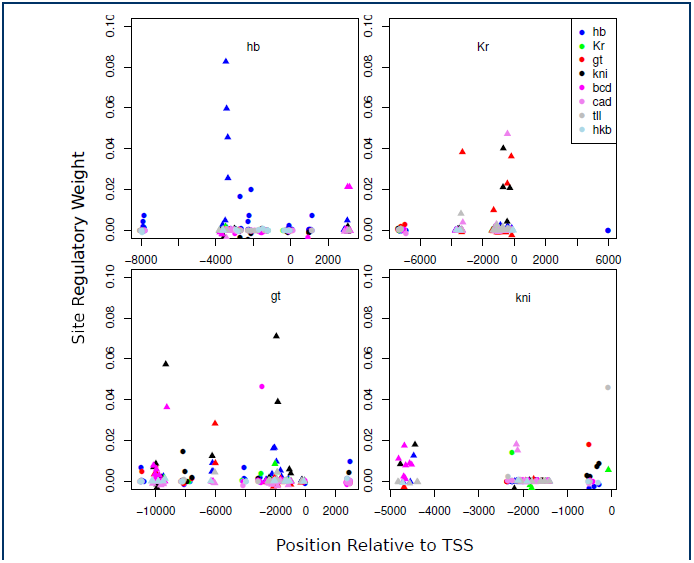
Plot of the regulatory weights of TFBS relative to their position in a regulatory region. The binding sites for different TF are shown in different color. The transcription start site is at zero position. Results for *hb* regulatory region are presented relative to TSS of the longest transcript. Sites within CRE are shown as triangles, sites outside CRE are drawn with circles.

Consequently, we arranged the sites in the order of increasing RWs and investigated how the removal of a different number of sites with the lowest RW influences the quality of model solution, which we evaluated by calculating the relative *RSS* score. As it is evident from Figure 7 the removal of as little as 10 TFBS with smallest RW results in 10% corruption of the model output. As a number of removed sites increases the model quality rapidly deteriorates. This *in silico* experiment demonstrates that sites with low RW are also important for the model output.

**Figure 7.**
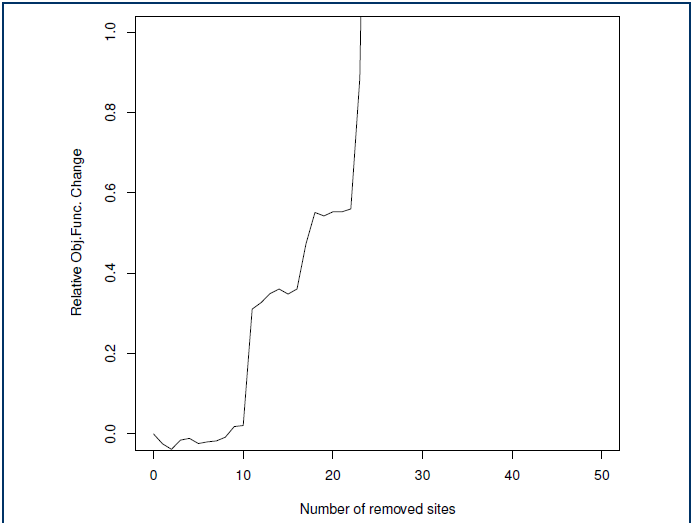
Impact of site removal on the quality of model output. The sites were removed in increasing order of their regulatory weights.

To study the spatial arrangement of the functionally important sites we constructed a new set of sites by filtering out the sites outside CREs (Table 5). We use this set and parameters obtained by fitting to the full set of TFBS to simulate gap gene expression patterns. As it is evident from Figure 8 the exclusion of sites located outside CREs worsens the quality of model output (*rms* = 34.28 as opposed to *rms* = 28.42 with full set of sites), but does not lead to the full pattern corruption.

**Table 5.**
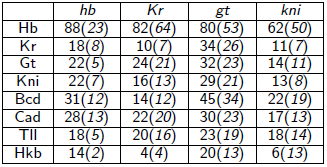
Total number of sites used in the model. Columns correspond to target genes, rows to TFs. The number of sites present in known CREs is given in brackets.

|  | *hb* | *Kr* | *gt* | *kni* |
| --- | --- | --- | --- | --- |
| Hb | 88(23) | 82(64) | 80(53) | 62(50) |
| Kr | 18(8) | 10(7) | 34(26) | 11(7) |
| Gt | 22(5) | 24(21) | 32(23) | 14(11) |
| Kni | 22(7) | 16(13) | 29(21) | 13(8) |
| Bcd | 31(12) | 14(12) | 45(34) | 22(19) |
| Cad | 28(13) | 22(20) | 30(23) | 17(13) |
| Tll | 18(5) | 20(16) | 23(19) | 18(14) |
| Hkb | 14(2) | 4(4) | 20(13) | 6(13) |

**Figure 8.**
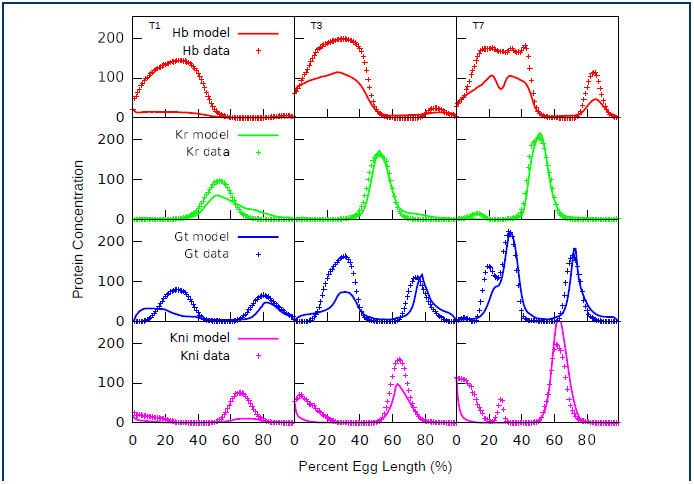
Model solution for sites within CREs as compared to gene expression data from the FlyEx database. The parameters were obtained by fitting to the full set of TFBS.

By visual examination of the plot (Figure 6) we selected the threshold value *w_r_* equal to 0.005 and further analyzed the sites with *w_r_* exceeding this threshold. The *hb* regulatory region contains 11 such sites for Hb and Bcd (see Table SI in Additional File 1). Two CREs are identified in this region. The *hb* anterior activator that is both necessary and sufficient for anterior *hb* expression is located about 200 bp upstream of the P2 promoter [39, 40] and contains several weak and strong binding sites for Bcd [39, 41] and Hb [42]. Late zygotic expression in the posterior cap and stripe, as well as PS4 is driven from both P1 and P2 promoters under control of the *hb* upstream enhancer located about 3 kb upstream of the P1 promoter [43, 44]. This element is regulated by Hb and has several predicted Tll and Kr TFBS [44, 43]. We found that 2 and 4 of 11 sites fall within anterior activator and upstream enhancer correspondingly (see Table S1 in Additional file 1). Interestingly both of the anterior activator sites overlap with strong Bcd sites annotated and verified by DNase I footprinting (Table 6).

**Table 6.**
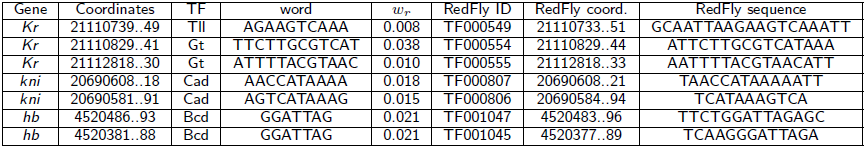
Sites with regulatory the weight *w_r_* > 0.005 that overlap with annotated DNase I footprint sites from RedFly database.

The *Kr* regulatory region contains 10 sites for Gt, Kni, Tll and Cad with RW exceeding the theshold. All this sites fall within different CREs contained in the region (see Table S2 of Additional File 1). Both Gt and Tll sites within the *Kr*_*730* CRE overlap with annotated DNase I footprint sites (Table 6). Both *Kr*_*730* and *Kr*_*CD1* elements produce *Kr* expression in the central domain [26, 27, 45, 46].

We identified 28 sites with RW exceeding the threshold for Bcd, Hb, Kr, Kni and Gt in the *gt* regulatory region. The (*gt*_*(-1)*) CRE drives *gt* expression in both anterior and posterior domains, while three other CREs reproduce reporter gene expression in the posterior (*gt*_*(-3)*) and different anterior domains (*gt*_*(-6*), *gt*_*(-10)*) [47, 36, 48]. Only 5 of identifed sites are located outside of CREs (see Table S3 of Additional file 1).

The *kni* regulatory region contains several CREs: *kni*_*(-5)* produces anterioven-tral expresssion, *kni*_*223+64* drives expression in the abdominal region and consists of two discrete sub-elements, *kni*_*(+1)* produces expression in both regions. In *kni*_*223+64* the 223-bp sub-element contains one Hb and six Cad TFBS and drives Cad-dependent reporter expression, while the 64-bp sub-element has six binding sites for Bed and mediates Bed-dependent expression in the anterior part of the embryo. Interestingly, the anterior expression of the 64-bp element becomes repressed when Hb binds to the 223-bp element [37]. We found 19 sites with RW exceeding threshold for Bcd, Hb, Cad, Kr, Kni and Tll in *kni* regulatory region (see Table S4 of Additional file 1). Only 6 of these sites are located outside the *kni* CREs. It is important to note that two sites within *kni*_*(223)* sub-element overlap with Cad annotated sites confirmed by DNase I footprint assays (Table 6).

## Conclusions

To model the regulatory mechanisms underlying the formation of gap gene expression domains we followed the formalism proposed in [49] and developed a two-layer model, in which firstly the activation level of each target gene in each embryo nucleus and at each time moment was calculated and at the next step mRNA and protein concentrations for this gene were computed. For calculation of the activation level we adapted **and modified** the thermodynamic approach in the form proposed in [4]. To calculate mRNA and protein concentrations we used differential equations. This innovative approach allowed us to connect the DNA-level information to the system dynamics and thus to overcome a serious limitation of the pure thermodynamic-based models which are static by their nature.

We further modified the method proposed in [49] by replacing the regulatory parameters *a_a_* and *β_r_* by the genetic inter-connectivity matrix *T^ab^* and introduced the delay parameter τ in our differential equations to account for the average time between events of transcription initiation and corresponding protein synthesis. This makes it possible to translate the **elementary** regulatory events at the DNA level to the **level** of gene interactions.

Our modeling approach has clear limitations. The promoter state is calculated by using methods of statistical thermodynamics, while the actual expression products result from this promotor state following the dynamics prescribed by the differential equations. This combination of intrinsically static and dynamical methods in one model is only consistent when there is an evident separation of the timescales of corresponding processes, the equilibration process of TF-DNA binding in the nucleus and the production process of transcribed mRNA and translated protein molecules. Taking into account the complex nature of transcription in eukaryotes, we believe that this assumption is a reasonable approximation for *Drosophila* genes. One indication for the length of transcription time specific for gap genes is in the fact that the gap gene expression products appear only in late cleavage cycles during the early *Drosophila* development partially because early cycles are too short for appropriate mRNA maturation [2]. On the other hand, the assumption about equilibrium states of the enhancer binding configurations is also only an approximation. There are clear data showing that such processes as nonspecific binding of TF to DNA and the facilitated diffusion of nonspecifically bound TF to a specific site play their role [50]. Despite the thermodynamic approach proved its efficiency in multiple applications, its proper extension for modeling more dynamic binding configurations seems promising.

The model takes as input the affinities of predicted TFBS together with spatial TF concentration data. The output of the model are spatial and temporal patterns of four gap genes *hb*, *Kr*, *gt*, and *kni* in the form of protein concentration profiles over about one and a half hour of development.

We used four-fold cross validation test and fitting to random dataset to validate the model and proved its sufficiency in data description. The identifiability analysis showed that most model parameters except of some parameters describing regulation by Tll, Hkb and Cad are well identifiable.

We demonstrated that our model is able to correctly predict expression patterns in *Kr* null mutants and five different reporter constructs from fits to wild-type data only. This provides an independent proof of model correctness and opens a way for its application for deciphering the mechanisms of transcriptional regulation and gene expression.

We used our model in two ways. Firstly, at the level of gene interactions we reconstructed the gap gene network topology and demonstrated that the basic features of this topology are in agreement with previous modeling results and data from literature [14].

Secondly, at the DNA level we studied the impact of individual TFBS on the model output. We measured this impact by calculating the site regulatory weight as a normalized difference between the residual sum of squares error for the set of all annotated sites and the set, from which the site of interest was left out. We found that the regulatory weights of TFBS show very weak correlation with their PWM score. This suggests that the influence of a TFBS on the phenotype is to a great extent explained not by the binding strength *per se* but by the way the binding sites are involved in the gene regulatory network. We also demonstrated that sites with low regulatory weight are important for the model output. This result corroborates the hypothesis about the biological significance of weak regulatory interactions [38]. Our *in silico* experiments also showed that functional important site are not exclusively located in CREs but are rather dispersed through regulatory region. It is of importance that some of the sites with high functional impact in *hb*, *Kr* and *kni* regulatory regions coincide with strong sites annotated and verified in Dnase I footprint assays.

## Methods

### Transcription factor and gene expression data

We used protein concentrations of transcription factors (referred to as TF in the text) Bcd, Cad, Hb, Gt, Kr, Kni, Tll and Hkb from FlyEx database (http://urchin.spbcas.ru, [5]) as inputs to the model. This database contains data on segmentation gene expression at the protein level and at discrete time points of cycle 13 and eight time classes (T1-T8) of cycle 14A. To estimate unknown parameters we used the expression patterns of gap genes *hb*, *Kr*, *kni* and *gt* from the same database. Model predictions were tested using gap gene expression patterns from *Kr^−^* embryos obtained from *Kr*^1^ loss-of-function allele [17], as well as reporter constructs driving reporter gene expression from the *Kr*_*CDl*, *Kr*_*730*, *gt*_*(-3)*, *kni*_*kd* and *kni*.*223+64* CREs (see REDFly database [35]).

### Sequence data

For each of four gap genes *hb*, *Kr*, *kni* and *gt* we predicted binding sites for Bcd, Cad, Hb, Gt, Kr, Kni, Tll and Hkb in the region spanning 12 Kbp upstream and 6Kbp downstream of the transcription start site. Transcription factor binding sites were predicted with position weight matrices (PWMs) [51], which were used to calculate the log-odds score of a site [52]. The PWMs were described in [53] and can be found at http://www.autosome.ru/iDMMPMM/ (see also the Additional file 1). The PWM thresholds were selected as in [54].

We took in the model the TFBS overlaping with the DNase I accessibility regions, which correspond to open chromatin. It was recently shown that in open chromatin regions predictions of transcription factor binding sites based on DNA sequence and in vitro protein-DNA affinities alone achieve good correlation with experimental measurements of *in vivo* binding [55]. The result of TFBS prediction, as well as positions of the DNase I accessibility regions and known CREs from the REDFly database are presented for *Kr* gene in Figure 9 and for all other genes in Additional File 1. The total number of TFBS for each TF and each gap gene considered in the model is shown in Table 5.

**Figure 9.**
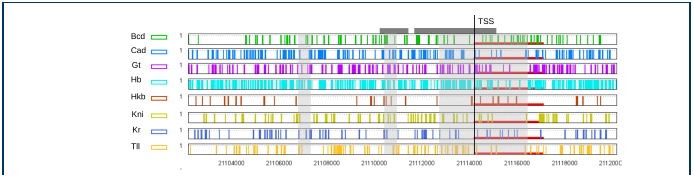
Prediction of binding sites in Kr’s regulatory region. The panels present predicted binding sites for eight TFs. The light-gray boxes denote the DNase accessibility regions, and the dark-gray bars mark positions of the RedFly CREs that drive gene expression in the blastoderm. The transcribed region of the locus is marked in red. Only the sites overlapping with the DNase accessibility regions were included in the model.

### The model

To model the regulatory mechanisms underlying the formation of gap gene expression domains we adapted the formalism proposed in [49] and developed a two-layer model, in which firstly the promoter occupancy (activation level) of each target gene in each embryo nucleus and at each time moment was calculated. The second layer of the model is based on differential equations and considers both mRNA and protein synthesis.

For calculation of promoter occupancy we adapted the thermodynamic approach in the form proposed in [4]:

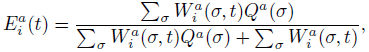

where *σ* is a molecular configuration of the regulatory region for gene *a*, *Q^a^*(*σ*) is the statistical weight of the interaction between TFs and bound basal transcriptional machinery (BTM), and 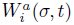(*σ,t*) is the statistical weight of configuration *σ* for nucleus-time coordinate (*i,t*), that depends on the concentration 
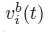
 of all TFs regulating gene *a* in nucleus *i* at time moment *t* (see [4] for details).

When cooperative binding is absent, we can write the statistical weight of a configuration *σ* as 
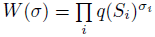
, where *σ*_*i*_ takes values 0 or 1 depending on whether site *S_i_* is occupied by its TF in the configuration or not. *q*(*S_i_*) is the strength of site *i* computed as

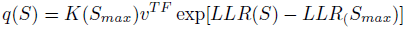

where *v^TF^* is TF concentration, *LLR*(·) is the log-odd score of a site, computed based on the known PWM of the TF and the background nucleotide distribution, *S_max_* is the strongest TFBS and *K*(*S_max_*) is its binding affinity constant.

In presence of cooperative binding, the statistical weight of a configuration is multiplied by a factor *ω*.

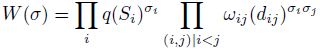

where *ω*_*ij*_(*d*_*ij*_) denotes the contribution to statistical weight due to interaction between the TFs bound to sites *S_i_* and *S_j_*, *ω*_*ij*_ is cooperativity constant, *d* represents the distance between the TF binding sites.

The statistical weight of the interaction between TF and bound BTM *Q*(*S*) is the product of the terms corresponding to each bound TF in the configuration. He and coathors [4] assumed that each TF is either an activator or repressor and proposed and “short-range” mechanism for repression based on the existing experimental work on a few well-characterized or synthetic CREs. A bound activator *A* interacts with the bound BTM with statistical weight *α_A_* > 1.

We assume that a bound repressor *R* does not directly interact with the BTM, but acts via short-range repression mechanism presumably making DNA in its “neighborhood” (defined by a range parameter *d_R_*) inaccessible to binding by any other TF. This configuration with statistical weight scaled by a factor of *β_R_* competes with those with the chromatin accessible to activators, thus effectively reducing the occupancy of activators. The parameter *β_R_* represents the strength of the repressor and may be interpreted as the equilibrium constant of the reaction that changes the chromatin state from accessible to inaccessible. There is no repression effect when *β_R_* is close to 0 while all activator sites are shut down in the neighborhood in the case of *β_R_* close to ∞.

In our model we consider that a TF can be activator for one target gene and repressor for another. Therefore we replaced the regulatory parameters *α_A_* and *β_R_* by the genetic inter-connectivity matrix *T^ab^*, which characterizes the action of TF *b* on gene *a*. The size of the matrix is *N_g_* × (*N_g_* + *N_e_*), where *N_g_* is the number of gap genes in the model (*N_g_* = 4) and *N_e_* is the number of external regulatory inputs (*bcd*, *cad*, *tll* and *hkb* genes, which are not regulated by gap genes, *N_e_* = 4). Consequently, *β_R_* is translated into negative components of the *T^ab^* matrix and positive ones become *α_A_* in order to calculate the activation level. This is done for each target gene in each nucleus and for each integration time step.

In the simplest approximation, the target gene expression level 
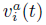
 is proportional to its activation level 
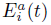
. We introduced the delay parameter τ to account for the average time between events of transcription initiation and corresponding protein synthesis, as the model is fitted to gene expression data at the protein level.

The second layer of the model is based on differential equations and considers both mRNA and protein synthesis. The equation for mRNA concentration 
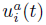
 of target gene *a* in nucleus *i* includes production, diffusion and decay terms, and the equation for protein concentration 
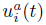
 describes protein synthesis, diffusion and degradation:

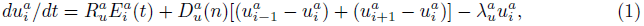

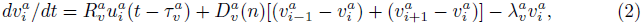

where *n* is the cleavage cycle number, 
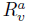
 and 
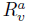
 are maximum synthesis rates, 
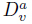
, 
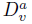
 are the diffusion coefficients, and 
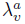
 and 
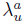
 are decay rates for protein and mRNA of gene *a*. The decay rates are related to the mRNA and protein half-lives 
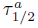
 by 
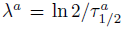
. The diffusion term was added to the equation for mRNA to smooth the resulting model output as it was too “spiky” without this term. The parameter 
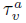
 is the delay parameter.

We model the dynamics of gap gene expression in cleavage cycles 13 and 14A and in one dimension along the central midline of the embryo. The cycle 14A is divided into eight temporal classes of 6.5 min each. The number of nuclei along the A-P axis is doubled when going from cycle 13 to 14A.

### Model fitting

The model was fitted to the protein concentration data for gap genes *hb*, *Kr*, *gt*, and *kni* from the FlyEx database. Parameter values were optimized by the differential evolution (DEEP) method, described in [18, 19].

The total number of optimized parameters in model (1)–(2) is 68. This includes 32 regulatory parameters *T^ab^*, 4 basal machinery constants, 8 binding affinity constants, 8 cooperativity constants, 4 range parameters for short range repression, 4 delay parameters and 8 decay rates. The diffusion constants and synthesis rates were fixed during the optimization.

We used the residual sum of squared differences between the model output and data (*RSS*) as the main objective function.

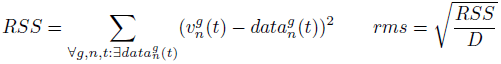

where *g, n* and *t* are gene, nucleus and time point respectively and *D* is the number of available experimental observations.

As was explained in [20], *RSS* can lead to counter-intuitive evaluations of the quality of fit and, therefore, we used the *weighted Pattern Generation Potential* proposed in this work as the second objective function:

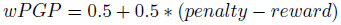

where

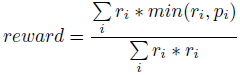

and

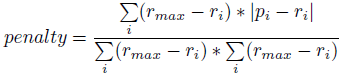

were *p_i_* and *r_i_* are respectively predicted and experimentally observed expression in nucleus *i*, *r_max_* is the maximum level of experimentally observed expression.

The third objective function penalizes the squared values of the regulatory parameters *T^ab^*:

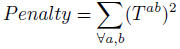

This function limits the growth of regulatory parameters, which have very wide ranges.

Consequently, the combined objective function is defined by:

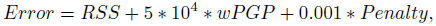

where the weights were obtained experimentally.

### Parameter identifiability

The parameter identifiability analysis was performed as described in [17]. The analysis finds non-identifiable parameters by calculating asymptotic confidence intervals [56, 57, 58]. The (1 – *α*)-confidence intervals for the parameter estimates *θ* are calculated in the vicinity of model solution as follows:

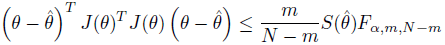

where the Jacobian *J*(*θ*) *= ∂RSS*(*θ*)/*∂θ* is the so called sensitivity matrix of size *N* × *m*, *F_α_*,*_m_*,*_N_*_*_m_* is an α-quantile of *F*-distribution with mand *N*-*m* degrees of freedom.

The confidence intervals of smaller size correspond to more reliable parameter estimates. In the case when the sign of the parameter estimate provides the most important feature, the estimate is assumed identifiable if the confidence interval is bounded away from zero.

The confidence intervals are overestimated for strongly correlated parameters. Correlation of parameters leads to computational errors since the sensitivity matrix is ill-conditioned.

Another method to study interrelations between parameters is the collinearity analysis [59]. The method is applied to reveal the so-called near collinear columns of the sensitivity matrix, namely the matrix of partial derivatives of the model solution with respect to the parameter vector. Identifiability of a parameter subset is estimated by collinearity index defined as

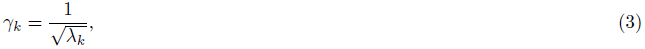

here λ_*k*_ is the minimal eigenvalue of the submatrix of the Fisher information matrix. High values of collinearity index means that the subset of parameters is poorly identifiable because at least two parameters in the subset are interrelated. More details on the methods can be found in [17].

## Competing interests

The authors declare that they have no competing interests.

## Author’s contributions

MS formulated the problem, MS, VG and KK formulated the model, MS, VG, IK and KK plan the experiments, KK and VG performed the experiments, IK calculated PWMs and discover the TFBSs, MS, VG, IK and KK interpret the results and wrote the paper.

## Acknowledgements

We are thankful to Prof Sergey Nuzhdin and Prof Alexander M.Samsonov for valuable discussions.The model derivation was supported by the RFBR grant 13-01-00405. Model fitting, validation, adaptation for studies of genetic variability, as well as analysis of model predictions and interpretation of results was done with support of the RSCF grant no.14-14-00302. The prediction of TFBS was supported by the “5-100-2020” Program of the Ministry of Education and Science of the Russian Federation. I.K. was supported by the Dynasty Foundation Fellowship.

## Additional Files

Additional file 1 — Supporting information

Positional weight matrices used to predict TFBS, positions of predicted binding sites in the regulatory regions of gap genes and lists of the binding sites with regulatory weight.

## References

1. Kaplan, T., Li, X.-Y., Sabo, P.J., Thomas, S., Stamatoyannopoulos, J.A., Biggin, M.D., Eisen, M.B.: Quantitative models of the mechanisms that control genome-wide patterns of transcription factor binding during early ¡italic¿drosophila¡/italic¿ development. PLoS Genet 7(2), 1001290 (2011).

2. Surkova, S., Kosman, D., Kozlov, K., Manu, Myasnikova, E., Samsonova, A., Spirov, A., Vanario-Alonso, C.E., Samsonova, M., Reinitz, J.: Characterization of the *Drosophila* segment determination morphome. Developmental Biology 313(2), 844–862 (2008)

3. Ay, A., Arnosti, D.N.: Mathematical modeling of gene expression: a guide for the perplexed biologist. Crit Rev Biochem Mol Biol 46(2), 137–151 (2011)

4. He, X., Samee, M.A.H., Blatti, C, Sinha, S.: Thermodynamics-based models of transcriptional regulation by enhancers: the roles of synergistic activation, cooperative binding and short-range repression. PLoS Comput. Biol. 6(9) (2010).

5. Pisarev, A., Poustelnikova, E., Samsonova, M., Reinitz, J.: FlyEx, the quantitative atlas on segmentation gene expression at cellular resolution. Nucleic Acids Research 37, 560–566 (2008). http://nar.oxfordjournals.org/cgi/reprint/gkn717v1.pdf

6. Simpson-Brose, M., Treisman, J., Desplan, C: Synergy between the Hunchback and Bicoid morphogens is required for anterior patterning in *Drosophila*. Cell 78, 855–865 (1994)

7. Arnosti, D., Gray, S., Barolo, S., Zhou, J., Levine, M.: The gap protein Knirps mediates both quenching and direct repression in the *Drosophila* embryo. The EMBO Journal 15, 3659–3666 (1996)

8. Gray, S., Levine, M.: Short-range transcriptional repressors mediate both quenching and direct repression within complex loci in *Drosophila*. Genes and Development 10, 700–710 (1996)

9. Hewitt, G.F., Strunk, B., Margulies, C, Priputin, T., Wang, X.D., Amey, R., Pabst, B., Kosman, D., Reinitz, J., Arnosti, D.N.: Transcriptional repression by the *Drosophila* Giant protein: Cis element positioning provides an alternative means of interpreting an effector gradient. Development 126, 1201–1210 (1999)

10. Lebrecht, D., Foehr, M., Smith, E., Lopes, F.J.P., Vanario-Alonso, C.E., Reinitz, J., Burz, D.S., Hanes, S.D.: Bicoid cooperative DNA binding is critical for embryonic patterning in *Drosophila*. Proceedings of the National Academy of Sciences USA 102, 13176–13181 (2005)

11. Fakhouri, W.D., Ay, A., Sayal, R., Dresch, J., Dayringer, E., Arnosti, D.N.: Deciphering a transcriptional regulatory code: modeling short-range repression in the Drosophila embryo. Molecular Systems Biology 6, 341 (2010)

12. Kulkarni, M.M., Arnosti, D.N.: cis-Regulatory logic of short-range transcriptional repression in *Drosophila melanogaster*. Molecular and Cellular Biology 25, 3411–3420 (2005)

13. Arnosti, D.N., Kulkarni, M.M.: Transcriptional enhancers: Intelligent enhanceosomes or flexible billboards?. Journal of Cellular Biochemistry 94(5), 890–898 (2005).

14. Jaeger, J.: The gap gene network. Cellular and Molecular Life Sciences 68, 243–274 (2011)

15. Reinitz, J., Sharp, D.H.: Mechanism of *eve* stripe formation. Mechanisms of Development 49, 133–158 (1995)

16. Jaeger, J., Surkova, S., Blagov, M., Janssens, H., Kosman, D., Kozlov, K.N., Manu, Myasnikova, E., Vanario-Alonso, C.E., Samsonova, M., Sharp, D.H., Reinitz, J.: Dynamic control of positional information in the early *Drosophila* embryo. Nature 430, 368–371 (2004)

17. Kozlov, K., Surkova, S., Myasnikova, E., Reinitz, J., Samsonova, M.: Modeling of gap gene expression in Drosophila Kruppel mutants. PLoS Comput. Biol. 8(8), 1002635 (2012).

18. Kozlov, K., Samsonov, A.: Deep - differential evolution entirely parallel method for gene regulatory networks. Journal of Supercomputing 57, 172–178 (2011)

19. Kozlov, K., Ivanisenko, N., Ivanisenko, V., Kolchanov, N., Samsonova, M., Samsonov, A.M.: Enhanced Differential Evolution Entirely Parallel Method for Biomedical Applications. LNCS 7979 V. Malyshkin (Ed.): PaCT 2013, 409–416 (2013)

20. Samee, M.A.H., Sinha, S.: Evaluating thermodynamic models of enhancer activity on cellular resolution gene expression data. Methods 62, 79–90 (2013)

21. Sachs, L: Applied Statistics. Springer, US (1982).

22. Dresch, J.M., Richards, M., Ay, A.: Thermodynamic modeling of transcription: sensitivity analysis differentiates biological mechanism from mathematical model-induced effects. BMC Systems Biology 4(142), 2–11 (2010)

23. Gaul, U., Jackle, H.: Pole region-dependent repression of the *Drosophila* gap gene *Krüppel* by maternal gene products. Cell 51, 549–555 (1987)

24. Hülskamp, M., Pfeifle, C, Tautz, D.: A morphogenetic gradient of Hunchback protein organizes the expression of the gap genes *Krüppel* and *knirps* in the early *Drosophila* embryo. Nature 346, 577–580 (1990)

25. Kraut, R., Levine, M.: Mutually repressive interactions between the gap genes *giant* and *Krüppel* define middle body regions of the *Drosophila* embryo. Development 111, 611–621 (1991)

26. Hoch, M., Schröder, C, Seifert, E., Jäckle, H.: Cis-acting control elements for *Krüppel* expression in the *Drosophila* embryo. The EMBO Journal 9, 2587–2595 (1990)

27. Hoch, M., Seifert, E., Jäckle, H.: Gene expression mediated by cis-acting sequences of the *Krüppel gene* in response to the *Drosophila* morphogens Bicoid and Hunchback. The EMBO Journal 10, 2267–2278 (1991)

28. Kerrigan, L.A., Croston, G.E., Lira, L.M., Kadonaga, J.T.: Sequence-specific transcriptional antirepression of the *Drosophila Krüppel gene* by the GAGA factor. The Journal of Biological Chemistry 266, 574–582 (1991)

29. Schulz, C, Tautz, D.: Autonomous concentration-dependent activation and repression of *Krüppel* by *hunchback* in the *Drosophila* embryo. Development 120, 3043–3049 (1994)

30. Casanova, J.: Pattern formation under the control of the terminal system in the *Drosophila* embryo. Development 110, 621–628 (1990)

31. Brönner, G., Jäckle, H.: Control and function of terminal gap gene activity in the posterior pole region of the *Drosophila* embryo. Mechanisms of Development 35, 205–211 (1991)

32. Rivera-Pomar, R., Niessing, D., Schmidt-Ott, U., Gehring, W.J., Jäckle, H.: RNA binding and translational suppression by Bicoid. Nature 379, 746–749 (1996)

33. Furriols, M., Casanova, J.: In and out of torso RTK signaling. The EMBO Journal 22, 1947–1952 (2003)

34. Dequeant, M.-L, Pourquie, O.: Segmental patterning of the vertebrate embryonic axis. Nat Rev Genet 9, 370–382 (2008)

35. Gallo, S.M., Gerrard, D.T., Miner, D., Simich, M., Des Soye, B., Bergman, CM., Halfon, M.S.: Redfly v3.0: Toward a comprehensive database of transcriptional regulatory elements in *Drosophila*. Nucleic Acids Research 10, 1–6 (2010)

36. Schroeder, M.D., Pearce, M., Fak, J., Fan, H.-Q., Unnerstall, U., Emberly, E., Rajewsky, N., Siggia, E.D., Gaul, U.: Transcriptional control in the segmentation gene network of *Drosophila*. PLoS Biology 2, 271 (2004)

37. Rivera-Pomar, R., Lu, X., Perrimon, N., Taubert, H., Jäckle, H.: Activation of posterior gap gene expression in the *Drosophila* blastoderm. Nature 376, 253–256 (1995)

38. Biggin, M.D.: Animal transcription networks as highly connected, quantitative continua. Developmental cell 21(4), 611–626 (2011)

39. Driever, W., Nüsslein-Volhard, C: The Bicoid protein is a positive regulator of *hunchback* transcription in the early *Drosophila* embryo. Nature 337, 138–143 (1989)

40. Struhl, G., Struhl, K., Macdonald, P.M.: The gradient morphogen Bicoid is a concentration-dependent transcriptional activator. Cell 57, 1259–1273 (1989)

41. Driever, W., Thoma, G., Nüsslein-Volhard, C: Determination of spatial domains of zygotic gene expression in the *Drosophila* embryo by the affinity of binding sites for the Bicoid morphogen. Nature 340, 363–367 (1989)

42. Treisman, J., Desplan, C: The products of the *Drosophila* gap genes *hunchback* and *Krüppel* bind to the *hunchback* promoters. Nature 341, 335–337 (1989)

43. Margolis, J.S., Borowsky, M.L., Steingrimsson, E., Shim, C.W., Lengyel, J.A., Posakony, J.W.: Posterior stripe expression of *hunchback* is driven from two promoters by a common enhancer element. Development 121, 3067–3077 (1995)

44. Lukowitz, W., Schröder, C, Glaser, G., Hülskamp, M., Tautz, D.: Regulatory and coding regions of the segmentation gene *hunchback* are functionally conserved between *Drosophila virilis* and *Drosophila* melanogaster. Mechanisms of Development 45, 105–115 (1994)

45. Hoch, M., Gerwin, N., Taubert, H., Jäckle, H.: Competition for overlapping sites in the regulatory region of the Drosophila gene Krüppel. Science 256, 94–97 (1992)

46. Capovilla, M., Eldon, E.D., Pirrotta, V.: The giant gene of Drosophila encodes a b-ZIP DNA-binding protein that regulates the expression of other segmentation gap genes. Development 114, 99–112 (1992)

47. Berman, B.P., Nibu, Y., Pfeiffer, B.D., Tomancak, P., Celniker, S.E., Levine, M., Rubin, G.M., Eisen, M.B.: Exploiting transcription factor binding site clustering to identify cis-regulatory modules involved in pattern formation in the Drosophila genome. Proceedings of the National Academy of Sciences USA 99, 757–762 (2002)

48. Segal, E., Raveh-Sadka, T., Schroeder, M., Unnerstall, U., Gaul, U.: Predicting expression patterns from regulatory sequence in Drosophila segmentation. Nature 451, 535–540 (2008)

49. Dresch, J.M., Thompson, M.A., Arnosti, D.N., Chiu, C.: Two-Layer Mathematical Modeling of Gene Expression: Incorporating DNA-Level Information and System Dynamics. SIAM J. APPL. M ATH. 73(2), 804–826 (2013)

50. Sharon, E., van Dijk, D., Kalma, Y., Keren, L., Manor, O., Yakhini, Z., Segal, E.: Probing the effect of promoters on noise in gene expression using thousands of designed sequences. Genome research, 168773–113 (2014).

51. Kulakovskiy, I.V., Makeev, V.J.: Discovery of dna motifs recognized by transcription factors through integration of different experimental sources. Biophysics 54(6), 667–674 (2009).

52. Lifanov, A.P., Makeev, V.J., Nazina, A.G., Papatsenko, D.A.: Homotypic regulatory clusters in drosophila. Genome Research 13(4), 579–588 (2003). http://genome.cshlp.org/content/13/4/579.full.pdf+html

53. Kulakovskiy, I.V., Boeva, V.A., Favorov, A.V., Makeev, V.J.: Deep and wide digging for binding motifs in ChIP-Seq data. Bioinformatics 26(20), 2622–3 (2010).

54. Kulakovskiy, I.V., Favorov, A.V., Makeev, V.J.: Motif discovery and motif finding from genome-mapped DNase footprint data. Bioinformatics 25(18), 2318–25 (2009).

55. Struffi, P., Corado, M., Kaplan, L., Yu, D., Rushlow, C., Small, S.: Combinatorial activation and concentration-dependent repression of the Drosophila even skipped stripe 3+7 enhancer. Development 138, 4291–4299 (2011)

56. Ashyraliyev, M., Jaeger, J., Blom, J.G.: On parameter estimation and determinability for drosophila gap gene circuits. BMC Systems Biology 2(83) (2008).

57. Bates, D.M., Watts, D.G.: Nonlinear Regression Analysis and Its Applications. JOHN WILEY & SONS, INC., New York, NY (1988)

58. Myasnikova, E., Kozlov, K.N.: Statistical method for estimation of the predictive power of a gene circuit model. J Bioinform Comput Biol 12(2), 1441002 (2014).

59. Brun, R., Reichert, P., Künsch, H.R.: Practical identifiability analysis of large environmental simulation models. Water Resour. Res. 37(4), 1015–1030 (2001).

